# AI-directed gene fusing prolongs the evolutionary half-life of synthetic gene circuits

**DOI:** 10.1101/2025.02.10.637495

**Authors:** Itamar Menuhin-Gruman, Matan Arbel, Doron Naki, Shaked Bergman, Tamir Tuller

## Abstract

Evolutionary instability is a persistent challenge in synthetic biology, often leading to the loss of heterologous gene expression over time. Here, we present STABLES, a novel gene fusion strategy that links a gene of interest (GOI) to an essential endogenous gene (EG), with a “leaky” stop codon in between. This ensures both selective pressure against deleterious mutations and high expression of the GOI. By leveraging a machine learning (ML) framework, we predict optimal GOI-EG pairs based on bioinformatic and biophysical features, identify linkers likely to minimize protein misfolding, and optimize DNA sequences for stability and expression. Experimental validation in *Saccharomyces cerevisiae* demonstrated significant improvements in stability and productivity for fluorescent proteins and human proinsulin. The results highlight a scalable, adaptable and organism-agnostic method to enhance the evolutionary stability of engineered strains, with broad implications for industrial biotechnology and synthetic biology.

## Introduction

Synthetic biology enables the engineering of biological systems for diverse applications, including therapeutic protein production, biosensing, and biomanufacturing ^1–14^. A key challenge in scaling these systems is maintaining the stability of engineered genes over evolutionary timescales ^15–30^. Heterologous gene expression often imposes a metabolic burden on host organisms, creating a selective advantage for mutants that reduce or eliminate expression. Over time, this leads to the loss of functionality and significantly impairs the viability of engineered systems for industrial or environmental use ^28,31–36^. In addition, this adds regulatory concerns and limits use of synthetic biology out of the lab, as it leads to lack of control over the generated sequences ^2,8,37–39^.

Several approaches have been explored to address evolutionary instability of the Gene Of Interest (GOI). Some strategies include generating libraries of complementary parts ^40,41^, limiting the user to selected elements and managing only the problem of repetitive elements – a partial solution. Others entail fine-tuning population dynamics ^23,42–44^, requiring designing several strains, tailored solutions, and much experimental tweaking. Many strategies attempt coupling gene expression of the GOI to the expression of an essential gene, and thus to host fitness. Some do so by engineering gene overlap ^15,45,46^ – a solution requiring much computational design, which works only for highly-specific cases where such overlap is possible. Others require designing biosensors which detect the protein of interest or byproducts, and activate essential genes based on these biosensors – this solution is highly specific and requires significant effort, if possible ^47,48^. Yet another solution suggests using the same promoter for both genes, on separate reading frames – mutations to the promoter which prove deleterious to the GOI, would also prove deleterious to the essential gene, leading to loss of fitness^22^. All these strategies have shown some success but are limited by technical complexity, preventing only specific mutation types, lack systematic tools, or are constrained to specific settings, genes or organisms.

In this paper, we introduce STABLES (Stop-codon Tunable Alternative Bifunctional mRNA Leading to Expression and Stability). It is a comprehensive approach to enhancing evolutionary stability through gene fusion. Our strategy involves physically linking the GOI to an EG via a shared promoter, on a single ORF, coupled with a “leaky” linker to enable differential expression levels ^49–52^. To optimize this system, we developed a machine learning (ML) tool that predicts the best EG partners for a given GOI, selects linkers likely to minimize protein misfolding using biophysical models of disorder ^53–60^, and generates codon-optimized DNA sequences for stability and high expression ^21,61–63^.

We validated STABLES in *Saccharomyces Cerevisiae* by stabilizing the expression of fluorescent proteins and the industrially relevant protein human proinsulin ^64^. GOI fused to selected EGs showed significantly enhanced stability and production over successive generations compared to controls. This study provides a scalable, flexible framework for stabilizing synthetic genes, offering a broad range of potential applications in synthetic biology and biomanufacturing.

## Results

### Overview of the STABLES Fusion Strategy

In this paper, we introduce STABLES, a comprehensive approach to enhancing evolutionary stability through gene fusion. It is host and GOI agnostic, robust to many mutations, and provides a generic, systematic, simple framework. Our design includes the following components (Fig. 1):

1. The gene of interest (GOI), to be expressed in the host organism.
2. An endogenous gene (EG), selected for optimal gene expression and mutational stability using an ML model. This model is based on meaningful bioinformatic features, and empirical data in S. Cerevisiae. The GOI and EG are expressed on a shared promoter, on a single ORF, where the GOI’s C-terminus is fused to the EG’s N-terminus.
3. The linker between them is selected for reducing likelihood of protein misfolding, using biophysical models of disorder ^53–60^.
4. The fusion gene is optimized for gene expression and avoidance of mutationally unstable sites^21,61–63^. This includes optimization of the GOI, linker, and depending on use case, the EG as well.
5. A leaky stop codon is placed after the GOI. This is a stop codon with a positive rate of read-through. This leads to the generation of two proteins – either the GOI alone, or the fusion protein. The rate of expression for the two proteins can be controlled, through selection of an appropriate rate of read-through ^49–52^. A codon is selected such that the fusion protein is produced in barely viable quantities for the host’s growth, while maintaining much higher expression in favor of the GOI’s protein alone - as this is the derived, relevant product. By ensuring just barely viable quantities of the fusion protein, the host’s mutational stability is further enhanced, as many more mutations would prove deleterious.
6. The EG in its native form is deleted from the host and replaced by this gene fusion. The host is now dependent on the fusion protein to provide the original EG function. Many mutations that would have reduced production or caused misfolding of the GOI, whether in GOI or promoter, would now reduce production of the fusion protein beneath viable quantities. Due to the host’s dependence on this protein, this leads to host lethality, and these mutations do not take hold in the population.

**Figure 1.**
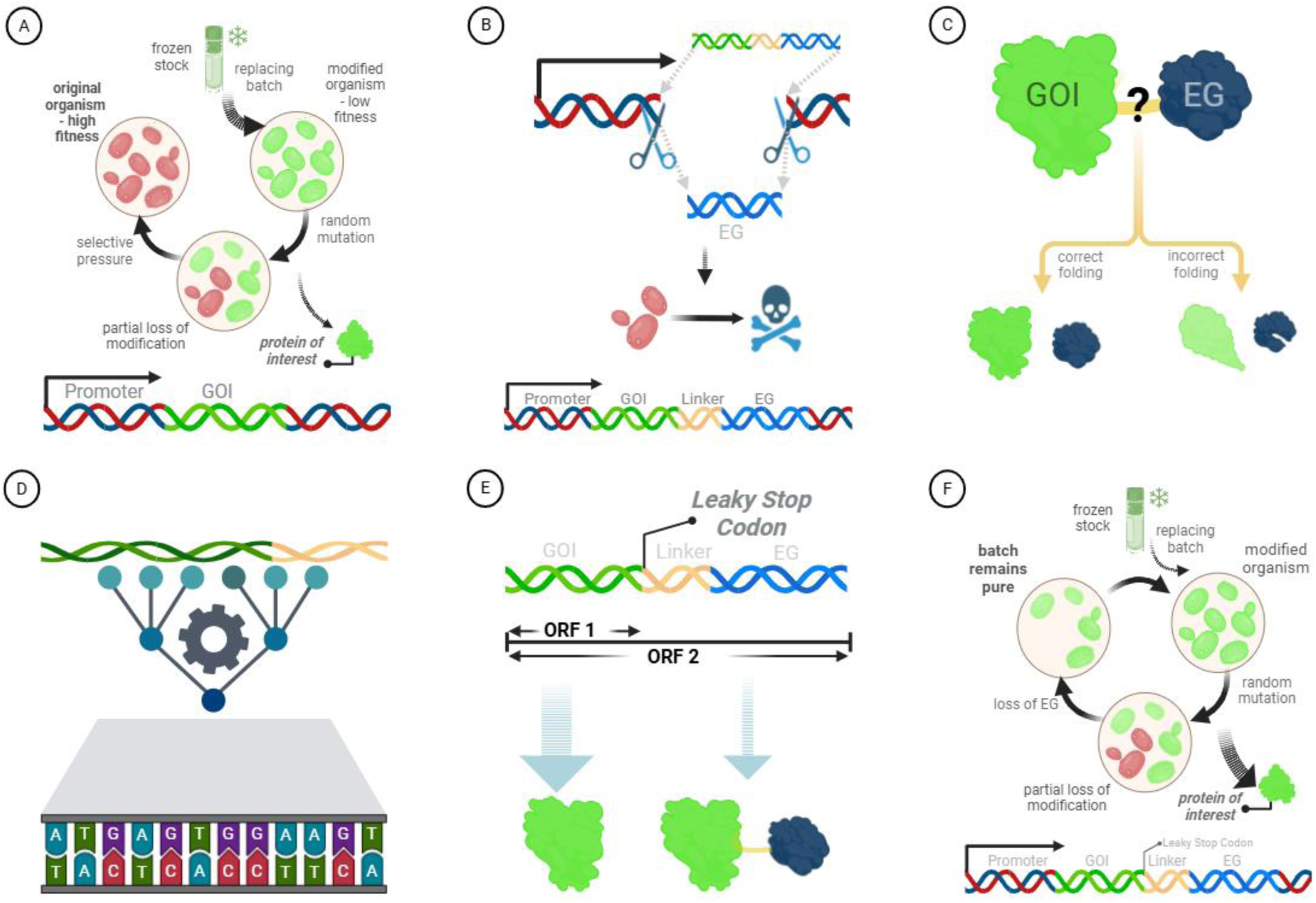
components of the STABLES solution: A. Standard production of a heterological gene. The heterologous Gene of Interest (GOI) is inserted into the genome. Any mutations that reduce expression or induce misfolding would prove advantageous due to the lower metabolic burden. These mutations proliferate, and a batch must be replaced. **B. Replacement of an endogenous gene (EG) with a fusion gene.** An endogenous gene is removed and replaced by a fusion gene composed of the GOI and EG. Mutations leading to loss of expression or misfolding would be deleterious to the EG, resulting in host death. This limits the spread of many mutations. The EG is selected by an ML model trained on experimental data. **C. Selection of linker.** Different linkers may lead to interaction between the fused proteins and misfolding. Using biophysical models and a database of fusion linkers, a linker is selected to minimize structural changes between the fused and unfused state. **D. Sequence optimization.** By optimizing the sequence of the GOI and linker, hypermutable sites are avoided, codon usage bias is maximized, and weak mRNA folding is enforced at the start of gene. This further improves stability and expression. **E. A leaky stop codon enables translation of both GOI and the fusion gene.** A leaky stop codon is placed between the GOI and linker. Due to partial read-through, both the protein of interest and the fusion protein are generated. By informed selection of stop codon, large quantities of the protein of interest and just viable quantities of the fusion protein are produced. The GOI’s mutational stability is further enhanced, as more mutations would prove deleterious. **F. Production of the heterological gene, aided by STABLES.** The new process has much higher mutational stability, reducing the need to replace batches. The optimized cells exhibit much higher expression.

### Fusion Strategy Improves Evolutionary Stability

To assess the impact of the fusion strategy, we evaluated the stability of GFP fused to various endogenous genes (EGs) by examining 10 strains from a previously described library of N-terminally GFP-tagged genes in *S. cerevisiae* ^65,66^. Fluorescence was used as a proxy for expression for 15 days.

This experiment was conducted prior to the creation of the EG selection mechanism – it was designed to give an indication of the need for such a model and its potential impact. For this purpose, meaningful bioinformatic features were generated for all EG, and they were clustered in many clustering configurations. 10 strains were selected such that they were consistently classified to different clusters and were near the centroids of these clusters. This generated a set of strains that were highly varied between them, while being representative of many similar genes. They were compared with a baseline strain of unfused GFP (see Methods section).

The experiment (Fig. 2.A.) validated the following conclusions, emphasizing the need for an EG selector:

1. Most strains demonstrated significant decline over the course of the experiment, emphasizing the existence of mutational instability and the need to improve it.
2. GOI-EG fusions exhibited significantly slower declines in fluorescence compared to unfused GFP, confirming that fusion of genes enhances stability.
3. Different EGs yielded varying degrees of stability, demonstrating that the stability of a fused gene depends on the endogenous gene selected.
4. One gene displayed a statistically significant advantage over unfused GFP, emphasizing the need for more informed gene selection.

**Figure 2.**
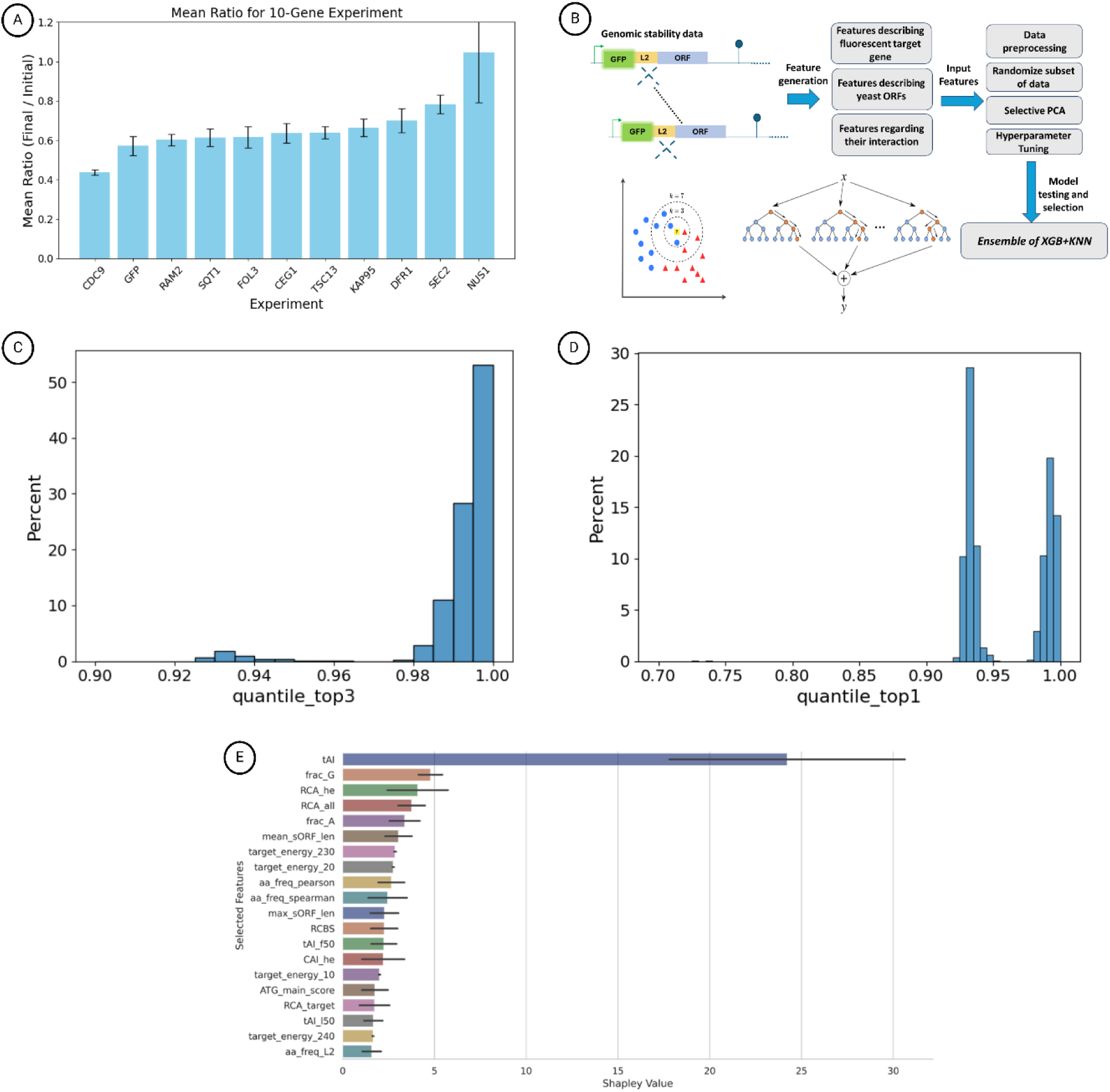
A computational model for EG selection: A. Motivation for developing EG selection model - Ratio between the final and initial fluorescence for 10 EG after 15 days. Each bar represents either GFP fused to a different endogenous gene, or the unfused GFP (2^nd^ from left). The value represents the mean ratio between final and initial fluorescence, for an experiment running for 15 days, with 3 repeats each. Fused genes maintain higher rates of fluorescence than unfused GFP at the end of the experiment (*student t* – *test*, *P* ≈ 0.048), thus more stable. The different experiments have different ratios throughout the experiment (*Kruskal* – *H*, *P* ≈ 0.035), demonstrating the need for gene selection. We have shown that SEC2 is more stable than unfused GFP (*student t* − *test*, *P* ≈ 0.047). We successfully predicted the top performing gene with our model. **B. Gene Selection Model Pipeline.** Experimental data ^65,66^ was used as input, comprising measurements for 6,685 EGs and 2 GOI. Biologically meaningful features were extracted to describe the EG, the GOI, and their interactions. For each randomized split of the training and validation data, feature processing and hyperparameter tuning were performed to optimize model performance. After analysing results across all splits, an ensemble approach combining K-Nearest Neighbours (KNN) and XGBoost (XGB) models was selected. **C. Performance among top 3 recommendations.** The different models were compared on their top recommendations. The test data was resampled with the bootstrap method, and the top recommendations were converted to quantile within this sample. The best performance among the top 3 candidates was presented (x-axis), indicating the expected performance when testing 3 fusion genes, and the height (y-axis) of each bin represents what percentile of samples fall within. Near-optimal performance is attained – a median score of 0.995. The probability of a score less than 0.98 is very low (*P* ≈ 0.048). **D. Performance for top recommendation.** A similar analysis was conducted, taking only the top candidate. Good performance can be expected, with a median score of 0.939. The probability of a score less than 0.92 is very low (*P* ≈ 0.007). **E. Distribution of Top 20 Shapley Values Across 20 XGB Models.** The graph shows the distribution of the top 20 Shapley Values, averaged across 20 XGB models, to assess feature importance in predicting fusion gene performance. The tRNA Adaptation Index (tAI) emerges as the most predictive feature. Following, features of importance include those related to GC content, codon usage bias, alternative ORF lengths, mRNA folding energy, and amino acid composition similarity between EG and GOI.

For statistical analysis, see supplementary 1.

### Machine Learning Predicts Optimal EG-GOI Combinations

The variability in stability observed across different EGs highlighted the importance of systematic EG selection. To address this, we developed a machine learning model to predict EG-GOI fusions that maximize expression and stability (Fig. 2.B., see Methods section). The model was trained on fluorescence data collected from GOI-EG fusion libraries under various conditions in *S. cerevisiae* (see supplementary 2) ^65,66^. As the fluorescence was measured after the variants had time to mutate, this is assumed to capture a combination of both expression and stability.

Using features such as codon usage bias (tRNA adaptation index ^67^, codon adaptation index ^68^), GC content ^69^, mRNA folding energy ^70,71^, ChimeraARS scores ^72^, and other meaningful bioinformatic features, the model successfully ranked potential fusion pairs.

In a reasonable use case, a user would generate and utilize very few genetic designs. The model would recommend 1-3 EG, which the user would validate experimentally, and proceed with the design exhibiting best performance. As the models were trained in cross-validation, each EG received a score equal to the quantile of its expression within the test set (e.g., 1.0 for EG with highest expression).Models were evaluated both on their expected performance (median score of EGs recommended by the model among cross-validations) and on their robustness (likelihood of recommending an EG with a low score). For performance measurement with more common and less relevant metrics, see supplementary.

Based on these evaluations, an ensemble model combining *k-nearest neighbours* ^73^ (KNN) and *XGBoost* ^74^ (XGB) was selected and trained. The KNN model exhibited a high median score, while the XGB model improved the model’s robustness. Selecting the best performance among top 3 candidates, the median score was 0.995, and the scores were above 0.98, (*P* < 0.05, Fig. 2.C.). When selecting only the top performer, the median score was 0.939, and the scores were above 0.92 (*P* < 0.001, Fig. 2.D.). These results underscore the predictive power of the ML model and its ability to systematically identify highly performing gene fusions, enhancing the efficiency of fusion design. Feature importance was calculated for the features, for further insight (Fig. 2.E.).

### Validation with Proinsulin Production

To demonstrate the real-world applicability of our approach, we applied it to stabilize human proinsulin expression in yeast, a biotechnologically relevant system. EGs were selected for high performance in both the XGB and KNN models, and additional engineering needs (see Methods). Thus, our model identified two EGs—CAF20 and ARC15—as suitable fusion partners for proinsulin.

A 30-day in-lab evolution experiment was conducted, where expression level measurements were taken every 5 days, using the ELISA protocol. The strains tested were the original proinsulin (as patented by Novo Nordisk), the proinsulin as optimized by the *Evolutionary Stability Optimizer* (ESO)^21^, and the optimized sequence fused to CAF20 and ARC15 as fusion genes (Fig. 3.A.). As in the 10-gene experiment, the unfused proinsulin replaced CAN1.

**Figure 3.**
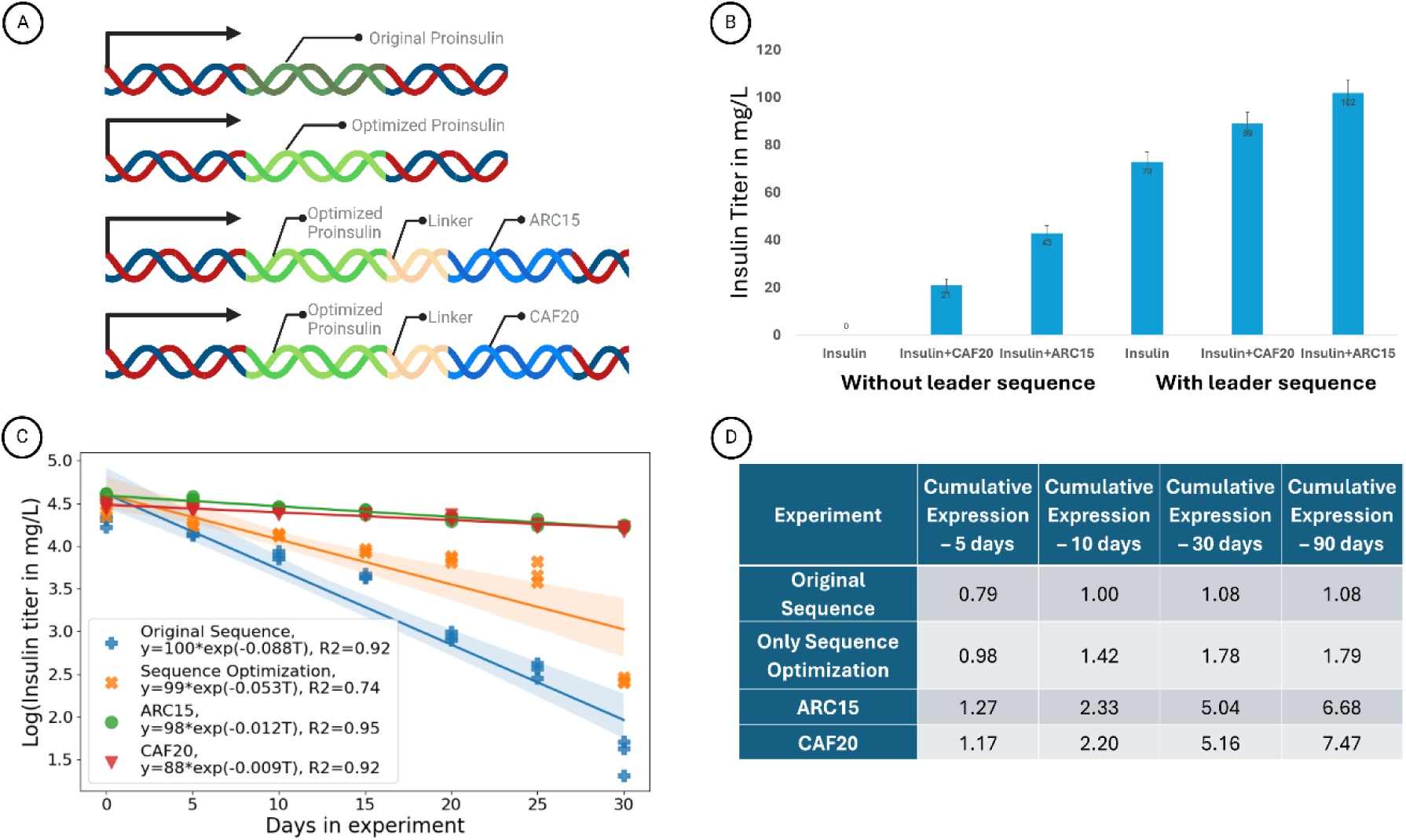
Demonstration of STABLES: improving expression of proinsulin in *S. cerevisiae*: **A. Design of the Proinsulin Evolution Experiment.** The design of the first proinsulin evolution experiment is displayed for clarity. **B. Gene fusion affects expression levels at time 0.** The expression level of proinsulin either unfused or fused to CAF20 or ARC15, with or without leader sequence, as measured by ELISA at the beginning of the lab evolution experiment. As previously demonstrated for the unfused case, the leader sequence is vital for expressing proinsulin. For the fused genes, it proved to be of high importance. This might be due to increased RNA stability or better localization of the protein in the cell. **C. Proinsulin production over time.** Proinsulin expression was measured by ELISA every 5 days, with 3 repeats, for each of 4 variants. The baseline variant is the original sequence, as patented by Novo Nordisk. The gene is placed instead of CAN1. The second variant is identical in structure, but with the proinsulin and linker optimized by ESO. The third and fourth are fusion genes, where proinsulin is fused with CAF20 and ARC15 respectively. The fusion genes were synthesized according to our design, including selection of EG, selection linker, addition of leader sequence, and sequence optimization. We assume exponential decay, supported by the high *R*^2^ on this linear estimation in log scale (*P* < 10^−6^). The curves have different decay rates (ANOVA F-test, *P* < 10^−19^). The variants of our design demonstrate a large improvement in stability. **D. Normalized Cumulative Proinsulin Generation Across Variants and Timespans.** Based on the exponential decay model, we can integrate the expression over time, giving us an estimate of the true value of interest - cumulative expression over time. We normalized all values by the estimated cumulative expression for the baseline variant over 10 days. This table presents the expected amount of proinsulin generated by each variant at different times. The variants of our design have significantly higher cumulative expression.

The need for a leader secretion sequence in proinsulin production in yeast has been well documented ^75–81^. However, it was not clear whether it may disrupt the efficacy of our design. We measured the expression at initiation for the original proinsulin and two fusion genes, with and without a leader sequence (Fig. 3.B.). All variants exhibited much higher expression in the presence of a leader sequence, and thus all further experiments were conducted as such.

The expression patterns approximately followed an exponential decay pattern, where the fused genes displayed a much slower decay rate (Fig. 3.C.). This enabled us to estimate the cumulative expression of proinsulin over time. These quantities were normalized by the expression estimated for 10 days, for the original proinsulin. The gene fusions showed a fivefold increase in total proinsulin yield over the experimental period (Fig. 3.D.). Interestingly, ARC15 demonstrated better performance for shorter durations (higher initial expression) while CAF20 demonstrated better performance for longer durations (slower decay) – the selection of the optimal gene will depend on the industrial use case.

For the fusion gene with ARC15, we measure 98.5 mg/L at initiation, where for the original sequence, we measure 72.9 mg/L – a ∼35% increase at initial expression. For the fusion gene with ARC15, we measure 68.7 mg/L after 30 days, meaning ∼70% of expression is maintained. For the original sequence, we measure 4.8 mg/L, meaning only ∼7% of expression is maintained.

Nanopore sequencing was conducted on the final sequences. It reveals that following the experiment, the fused proinsulin suffered few mutations, while the unfused proinsulin sequence was lost completely (Fig. 4.A.) ^82–85^. This has been further validated by western blot (Fig. 4.B.).

**Figure 4.**
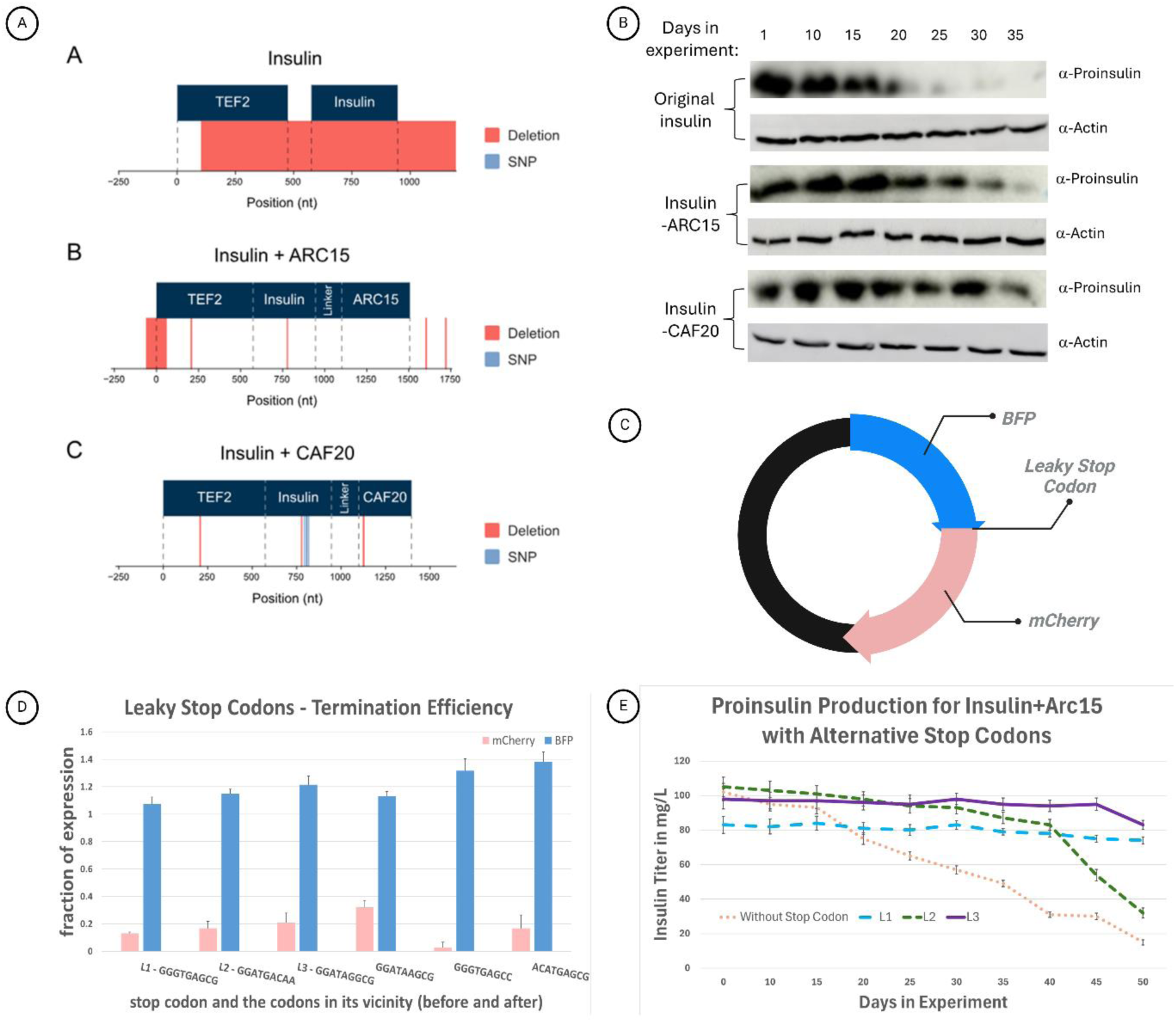
Experimental study of STABLES results and components: A. Mutation accumulated in an in-lab evolution experiment. ^88,89^. (A) Nano-pore sequencing of unfused proinsulin shows that after the in-lab evolution experiment, most of the promotor and the reading frame of insulin were erased from the plasmid. (B) Nano-pore sequencing of proinsulin fused to ARC15 reveals a significant deletion in the promoter, and a few small deletions in the promotor region and the proinsulin coding region. (C) Nano-pore sequencing of proinsulin fused to CAF20 shows a few mutations in the proinsulin coding region, the promotor and in CAF20. **B. Western Blot Validation.** Western Blot analysis validates our findings – the fused variants expressed proinsulin for much longer than the unfused variant. Actin measurements were taken for control. **C. Design of leaky stop codon construct.** The design of the stop codon experiment is displayed for clarity. **D. Determining the termination efficiency of different leaky stop codons.** BFP and mCherry were fused using different leaky stop codons in the linker. The fluorescence of each was measured as proxy for expression level, normalized by the non-fused expression level. The red bar is mCherry expression, which was fused to the C’ terminal of BFP, analogous to EG in our design. The blue bar is the expression of BFP, analogous to the GOI. In the graph, 3 representatives of the L1-3 stop codon design (see Fig. 4.E.) are presented, as are 3 representatives of codons not selected, due to read-through rate which was either too low or high. **E. Proinsulin concentration over time for different leaky stop codons.** Proinsulin was fused with ARC15 four times, each with a different stop codon. This graph illustrates the proinsulin concentration measured by ELISA over 50 days. Using any leaky stop codon improved the stability of the gene fusion (student t-test at t=40, *P* < 10^−5^). Different codons lead to different expression profiles (Kruskal H-test, *P* < 0.003), emphasizing the need for informed selection. The leaky stop codons were selected such that the last amino acid of proinsulin is maintained, we utilize the design principles outlined in ^86^, and select stop codons which displayed a read-through of 0.1-0.25 in the previous experiment. The sequences used for the stop codon and the 3 nt afterwards were: L1 – TAGGCG. L2 – TGAGCG. L3 – TGACAA.

### Leaky Stop Codon Enables GOI Production and Overexpression

To enable generation of both the fusion protein and the GOI alone, we utilized leaky stop codons, which allow partial translational read-through ^49–52^.

The ability to enforce low but non-zero read-through rates is a key factor in improving the stability and industrial viability of our design. If the fusion protein’s abundance is reduced to barely viable, then it is highly likely that any mutations affecting the GOI’s expression would lead to host lethality, promoting mutational stability. This is in tandem with the enforcement of much higher expression of the isolated GOI, which is the necessary and desired product.

Thus, informed selection of the stop codon is necessary. To enable this informed decision, we conducted an experiment, creating a fusion protein of BFP and mCherry, with different leaky stop codons between. This was expressed on a plasmid (Fig. 4.C.). The red fluorescence is expressed only in the fusion protein, while the blue fluorescence is expressed in both the fusion protein and BFP alone. Thus, the ratio between the fluorescence measurements can be used to derive the rate of read-through (Fig. 4.D.).

We selected the 3 stop codon designs expressing the lowest mCherry fluorescence (to minimize read-through) which was still detectable (to increase cell viability). We conducted another evolution expression, attaching the proinsulin gene to ARC15, either without a stop codon or with one of the 3 selected. This is in accordance with the STABLES design. Using a similar protocol, expression of proinsulin was measured over 50 days, this time capturing the presence of both the fusion protein and the isolated proinsulin. In addition to the secretion of isolated proinsulin (which is a requirement in and of itself), we observed a much higher expression rate, and cumulative expression accordingly (Fig. 4.E.).

For the fusion gene without a stop codon, we measure 102 mg/L at initiation, where for the best candidate L3, we measure 98 mg/L– a ∼4% decrease at initial expression. For the best candidate L3, we measure 83 mg/L after 50 days, meaning ∼85% of expression is maintained. For the fusion gene without a stop codon, we measure 15 mg/L, meaning only ∼15% of expression is maintained.

## Discussion

### An organism-agnostic strategy to tackle evolutionary instability

Previous studies proposed methods for increasing mutational stability of the GOI. Some of these methods are either dependent on host-specific pathways, or require design and implementation of complex parts (e.g. biosensors, overlapping genes) which may not be possible. Others require developing multiple strains and managing population dynamics, or provide robustness only to specific mutation types (e.g. repetitive parts, mutations in promoter). This study introduces STABLES, a simple, robust, generic, host and GOI-agnostic strategy to address evolutionary instability in synthetic biology. By fusing the GOI to an EG on the same ORF under a shared promoter, many deleterious mutations would lead to host lethality, leading to higher GOI mutational stability. By incorporating a leaky stop codon, we enable these gene fusions to generate high expression of the GOI alone, while increasing their mutational stability. These, together with informed selection of the linker, sequence optimization and use of a leader sequence, are key in the improved stability and performance observed in our experiments.

We find that following all our design principles, we observe a 15% decline in expression of proinsulin over 50 days. This contrasts with the original design, which declined by 93% over 30 days. This is in addition to a ∼30% increase in initial expression.

Our machine learning tool enhances the utility of this approach by selecting better GOI-EG pairings based on biologically meaningful features. Translational efficiency metrics, such as tRNA Adaptation Index (tAI), emerged as dominant predictors. A possible explanation for this result may be related to the fact that higher translation efficiency tends to be associated with genes that are highly expressed, have higher mRNA levels (due to higher mRNA stability, among other factors), and that are more conserved and thus tend to fold more efficiently ^87–91^. Other features, including GC content, RNA folding energy, alternative shifted ORFs, and amino acid composition that are probably also associated with higher expression, mRNA stability and robust protein folding.

Protein disorder predictions were employed to guide linker selection, increasing the likelihood that the gene fusions maintain native protein folding and functionality (See Methods). Combined with avoidance of mutational hotspots and sequence optimization for expression and stability, these elements ensure the practical applicability of the approach across a broad range of use cases.

### Future Directions

Using our machine learning model, we have already seen empirical evidence for good performance in predicting high-performing GOI-EG pairs for different GOI, as shown in our experimental validation. Its reliance on sequence-based features enables effective application to non-model organisms, even in the absence of extensive empirical data. However, expanding the dataset used for training the model remains a valuable avenue for further enhancement. Systematic testing for libraries from more host organisms and a broader range of target genes would improve the model’s generalizability and ensure its utility across an even wider variety of synthetic biology applications. Furthermore, in many organisms, epigenetic silencing must be considered when optimizing expression and stability ^24,92–99^. While we have designed our model with capabilities to avoid epigenetically silencing motifs, this application should be tested in vivo.

While our method has shown significant promise, further experimental validation is needed to refine certain aspects. For example, empirical testing of linker selection will help confirm computational predictions and guide future improvements. An in-depth analysis and testing of the leaky stop codons would also enable further optimization.

### Broader Implications

This study demonstrates the potential to stabilize synthetic genes in both industrial and environmental contexts. In biomanufacturing, the ability to maintain stable GOI expression over extended periods can reduce costs, improve scalability, and simplify regulatory processes. In environmental applications, robust gene fusions could support long-term deployment in dynamic and uncontrolled conditions, enabling breakthroughs in bioremediation and biosensing.

### Conclusion

By integrating gene fusion, machine learning-guided gene selection, protein disorder predictions for minimizing misfolding, sequence optimization, and leaky stop codons, we present a robust framework for addressing the evolutionary instability of synthetic genes. This system achieves high GOI expression, ensures host viability, and enhances mutational stability across generations. The reliance on biologically interpretable features, such as tAI and RNA folding energy, underscores the model’s capacity to identify high-performing designs.

Its organism-agnostic nature and focus on sequence-level optimization make this approach a versatile solution for diverse synthetic biology applications. With further validation, refinement, and the incorporation of additional data, this framework holds significant promise for advancing biotechnology and synthetic biology.

## Methods

### Overview of Gene Fusion Design

We developed a multi-step workflow to design and optimize GOI-EG fusion genes for enhanced evolutionary stability and expression. This process included the selection of EGs using an ML model, the identification of optimal linkers to minimize misfolding, sequence optimization of the synthetic gene to maximize expression and stability, and addition of a low-readthrough leaky stop codon.

### Machine Learning Model for EG Selection

The ML model was trained on fluorescence datasets from yeast libraries (see supplementary 1) ^65,66^. The dataset included GOI-EG fusion genes for 5,185 EG (∼78% of all EG in yeast), fused with either GFP or mCherry. Fluorescence was measured across multiple time points.

### Features and Model Architecture

The model utilized a diverse set of bioinformatic features derived from the sequences of each GOI and EG. Key feature families included:

1. **Codon Usage Bias:**

- **tRNA Adaptation Index (tAI):** Calculated for the full sequence, first 17 amino acids, and sliding windows across the GOI and EG ^67^.
- **Relative Codon Adaptation (RCA):** Evaluated for the GOI, EG, and sliding windows ^100^.
- **Codon Adaptation Index (CAI):** Assessed for the full sequence and sliding windows ^68^.
- **Effective Number of Codons (ENC):** Measures diversity in codon usage ^101^.
2. **Sequence Composition:**

- **GC Content:** Calculated globally and in sliding windows for both GOI and EG ^69^.
- **k-mer Frequencies:** Counts of nucleotide or amino acid substrings of lengths 3–5.
- **Amino Acid Frequency Correlation:** Spearman correlation between GOI and EG amino acid compositions.
3. **Thermodynamic Properties:**

- **Local Folding Energy:** Calculated using ViennaRNA ^102^ in sliding windows (50 bp).
4. **Chemical Properties:**

- **Molecular Weight and Hydrophobicity:** Computed for both GOI and EG sequences ^103^.
- **Isoelectric Point (pI):** Assessed at physiological pH.
5. **Translation Context:**

1. **Start Codon Context:** Features describing the nucleotide flanking regions of start codons.
2. **Shifted Open Reading Frame (sORF) Length:** Alternative reading frame lengths.
6. **ChimeraARS Score:**

- The ChimeraARS score quantifies sequence similarity to a reference set of genes with high codon usage bias. This metric enhances predictions of sequence stability beyond standard codon usage features ^72^.

The ML pipeline employed an ensemble model combining k-nearest neighbors (KNN) and XGBoost architectures (Fig. 3.A.). The KNN model emphasized features of the EG and provided high median performance - but was prone to failures. The XGB model emphasized features of the GOI, EG, and interaction features, and provided much higher robustness to failure. Hyperparameter tuning and cross-validation were conducted using Optuna to ensure robust performance (Details in supplementary).

### Feature Importance in Machine Learning Predictions

Feature importance was analyzed using SHAP ^104^ (SHapley Additive exPlanations) for XGBoost and forward feature selection for KNN. SHAP provided a global ranking of feature contributions, while forward selection iteratively identified the most impactful features for KNN predictions. The XGBoost results, which were more robust, provided the primary conclusions, while KNN analysis offered additional insights where consistent or complementary.

### XGBoost Feature Importance

The SHAP analysis (Figure 3.D.) revealed **tRNA Adaptation Index (tAI)** ^67^ as the dominant feature, significantly outpacing all others. Variants with high tAI scores consistently showed enhanced stability and expression, underscoring its central role in optimizing translation.

Following tAI, the next most important features were **GC content** ^69^ and **Relative Codon Adaptation (RCA)** ^100^, both reflecting translational optimization and sequence stability. Additional influential features included:

- **Amino Acid Frequency Correlation** between the GOI and EG, suggesting reduced metabolic burden for similar protein compositions.
- **Local Folding Energy** ^102^ downstream in the GOI and near the initiation site, emphasizing the importance of avoiding unstable mRNA secondary structures.
- **Shifted ORF Length** ^105–107^, **Ribosomal Coverage Bias Score (RCBS)** ^108^, and **tAI in the First 17 Amino Acids** ^109–112^, which further highlighted the significance of efficient translation, and successful initiation.

### KNN Feature Importance

The KNN forward selection supported the importance of codon usage-related features, particularly in the EG. Key features included **RCA**, **tAI**, and **CAI** ^68^ metrics within the EG and in specific regions, such as the first 17 amino acids. These findings complemented the more GOI-focused insights from XGBoost by emphasizing the role of the EG in overall gene fusion stability.

### Linkers for Protein Fusion

Protein misfolding remains a challenge in gene fusion strategies ^113–116^. While tools such as AlphaFold ^117^ offer accurate prediction of protein structure, they are too computationally intensive to enable a reasonable comparison between many linkers. Rather, we used tools such as IUPred2A ^71^, a biophysical model, and MoreRONN ^70^, a machine learning model, to predict protein disorder profiles and assess the impact of linkers. As protein disorder profiles have a profound effect of protein folding, it is taken as a proxy variable – linkers with smaller effect on the disorder profile are less likely to influence the protein folding, and thus less likely to cause misfolding ^57,58,118,119^. Linkers were selected to minimize disruptions to the disorder profile upon fusion, likely preserving native folding patterns of both the GOI and the EG ^53–59^. This minimization was applied by calculating the Euclidean distance between the disorder profiles before and after fusion, for the 1,280 linkers in a linker database ^59^. The linker inducing the smallest distance was selected.

Although experimental validation of the linker selection step remains pending, these predictions provide an essential foundation for mitigating misfolding risks in future applications.

### Scoring and Selection

Linkers were scored based on the L2 distance between disorder profiles of the unfused and fused states. The optimal linker was selected to minimize disruptions to native folding patterns, preserving the functionality of both the GOI and EG.

### Optimizing DNA Sequences for Stability and Expression

Sequence optimization was performed to enhance the stability and expression of the GOI-EG fusion genes using the **Evolutionary Stability Optimizer (ESO)** ^21^ pipeline. **ESO** integrates **DNAChisel** ^120^, a tool that optimizes genetic sequences while maintaining biological constraints. ESO has been previously validated for its ability to improve the evolutionary stability of synthetic genes. The ESO pipeline focuses on optimizing codon usage, minimizing mRNA instability, and ensuring the long-term expression of engineered genes by accounting for evolutionary pressures.

### Optimization Objectives

1. **mRNA Folding Optimization**: It has been demonstrated that minimizing the secondary structure formation in mRNA near the translation initiation site, maximizes the expression ^102,121–126^. The **local folding energy** of the first 15 codons in the mRNA sequence was calculated using ViennaRNA ^102^ and codons were selected to optimize weak folding energy.
2. **Codon Usage Optimization**: For the rest of the sequence, codon usage was optimized to match the tRNA abundances of *S. cerevisiae*. This was achieved by calculating and optimizing the **tRNA Adaptation Index (tAI)** ^67^ for the GOI, linker, and (depending on the application and relevancy) the EG, ensuring that codon usage patterns align with the host’s translational machinery.
3. **Stability Enhancements**: Sequences were optimized to avoid hypermutable regions ^24,93,127–136^. These optimizations ensure synthetic genes are tailored for host-specific transcription and translation efficiencies, enhancing long-term performance in diverse applications.

### Experimental Validation

#### Fusion Stability in *S. cerevisiae*

We validated the gene fusions by testing their ability to stabilize GOI expression in *S. cerevisiae*. Taking the Schuldiner lab library ^65,66^ (N’ SWAp Tag (SWAT)-GFP, derived from *S. cerevisiae* BY4741 background strain), we selected variants from it such that the EG are varied and representative. This was performed by performing k-means for *k* = 2 … 130, 5 iterations each, and selecting the 10 genes most consistently in different clusters, while closest to cluster center. As a control, they were compared with unfused GFP, integrated into the genome by replacing CAN1. The CAN1 gene encodes an arginine permease, which is not essential for yeast survival under laboratory conditions where arginine is supplemented in the medium ^137–141^.

Their yield was measured over 15 days of in-lab evolution, with 3 repeats each. The evolution experiment took place in 96-Deepwell Plates (Starlab group Cat.No S1896-2110). Fluorescence levels were taken as proxy for protein expression.

We first normalized this value by the OD, to control for yeast density. Furthermore, we performed this analysis for the wild type (WT) yeast, to find the inherent fluorescence of yeast. Following, we detracted this value from other measurements, to isolate the fluorescence due to GFP. ^68–72,140^

#### Proinsulin Production in Yeast

To test a biotechnologically relevant application, we applied the workflow to human proinsulin. The proinsulin sequence was obtained from Novo Nordisk patented sequence for proinsulin manufacturing in yeast (patent WO2014195452A1). EGs were selected using the following criteria:

1. EG was predicted to be in top 5% for XGBoost architecture.
2. EG was predicted to be in top 10% for KNN architecture, due to less robustness.
3. EG length less than 500nt, to ensure reasonable synthesis of the combined fusion protein.
4. EG has no internal repeats of length > 10, to ensure greater stability.
5. EG is not ribosomal, due to many interactions of ribosomal proteins, with many points of failure.
6. EG does not have induced function – they are more sensitive to changes in expression.
7. We chose genes from multiple cellular processes to decrease redundancy.

Based on these criteria, CAF20 and ARC15 were selected.

4 variants were compared:

1. The original proinsulin sequence, expressed on a high copy number plasmid (Prs426)
2. A similar variant, where the proinsulin sequence was optimized by the ESO ^21^.
3. A fusion gene composed of optimized proinsulin and CAF20 expressed on a high copy number plasmid in a strain that CAF20 was deleted from the genome.
4. A similar variant for ARC15.

Each variant included an alpha leader. The linker **1fzbf** was selected.

They were tested for yield over 35 days of in-lab evolution, with 3 repeats each. Proinsulin levels were quantified using ELISA plates according to manufacturer instructions (product number RAB0327) for proinsulin detection and verified by western blot using proinsulin specific antibody (sigma i2018).

The constructs were analyzed through Nanopore sequencing before and after the evolution experiment, for deeper understanding of the accumulated mutations. Nanopore reads were first processed using chopper ^82^, trimming the first 10nt of each read and keeping reads of length above 500. The reads were mapped with minimap2 ^83^ to a reference sequence, including the yeast genome and the relevant construct. Quantification of optimized variants’ mRNA levels was conducted using Salmon ^84^, and variant calling was conducted by DeepVariant ^85^.

#### Leaky Stop Codon Rate of Read-through

Selecting and implementing a leaky stop codon is required for our design. We require a rate of read-through such that:

1. The expression of the fusion protein is high enough to generate viable quantities of the fusion protein, required for the cell’s growth.
2. It is the lowest value reasonable, to prove more mutations deleterious.
3. High quantities of the GOI alone, as this maximizes the output of the system.

To enable an informed selection for our stop codons, we generated different stop codons using the design principles outlined in (Mangkalaphiban et al., 2021) ^86^. In this paper, Riboseq is conducted on all genes in S. Cerevisiae. They created a predictive model for rate of read-through and found the most influential feature to be the context around the stop codon – from 3 nts before to 6 after, 12 nts total. Further refining the model, they created a ranking of leakiness based on nt at each position. Using these rankings, we generated sequences predicted to be with high read-through rates.

We conducted an experiment where we implemented a fusion gene on a plasmid. This fusion gene was composed of BFP connected to the C’ terminus of mCherry, with the different leaky stop codons generated. The fluorescence of BFP is proxy for the expression of both the GOI and the fusion protein, and the fluorescence of mCherry is proxy for the expression of only the fusion protein.

We selected 3 stop codon designs such that:

1. The rate of read-through, as measured in the mCherry relative fluorescence, is in the range of 0.1-0.25.
2. The relative fluorescence of BFP, indicative of the expression of isolated GOI, is greater than 0.9.
3. The last amino acid in proinsulin is unchanged.

We conducted another evolution experiment. 4 variants were generated. In each, they followed the design of the ARC15 variant from the previous experiment. In one there was no stop codon, the other 3 followed the stop codon designs selected.

They were tested for yield over 50 days of in-lab evolution, with 3 repeats each. Proinsulin levels were quantified using ELISA plates according to manufacturer instructions (product number RAB0327) for proinsulin detection.

## Acknowledgements

S.B. is supported by a fellowship from the Edmond J. Safra Center for Bioinformatics at Tel Aviv University. We thank Prof. Maya Schuldiner lab, for providing us with SWAp-Tag strains and data. We thank Dr. Daniel Dovrat for helpful discussions and comments.

