## Supplementary information for "AI-directed gene fusing prolongs the evolutionary half-life of synthetic gene circuits"

**Supplementary 1: Statistical Analysis**

*Comparison of unfused to fused GFP*

In the 10-gene experiment (Fig. 3.1), we compared the stability of the 10 fused genes to the unfused GFP baseline. The error analysis process was as follows:

1. **Data Collection**:
   For the 10 fused genes, we calculated the mean (μ) and standard deviation (σ) of their fluorescence measurements at $t = 15$, the last day of the experiment. The fluorescence measurements were normalized by the fluorescence at $t = 1$, the experiment initiation. This was done in order be able to assume normal distribution, while maximizing the effect of decay. For the unfused GFP, we used the standard error (SE) of its fluorescence measurements at this time.

2. **Calculation of T-Score**:
   To determine if the fused genes were significantly more stable than the unfused GFP, we calculated a t-score for the mean fluorescence of the 10 fused genes using the formula:

$$t = \frac{\mu_{coupled} - \mu_{\text{unfused}}}{\sqrt{\frac{\Sigma_i {\sigma_i}_{coupled}^2}{n_{coupled}} + SE_{\text{unfused}}^2}}$$

   Where:

- $\mu_{\text{fused}}$ is the mean fluorescence for each of the fused genes.
- $\mu_{\text{unfused}}$ is the mean fluorescence of the unfused GFP.
- ${\sigma_i}_{\text{fused}}$ is the standard deviation of fluorescence measurements for the fused genes.
- $n_{\text{fused}}$ is the sample size for the fused genes.
- $SE_{\text{unfused}}$ is the standard error of the unfused GFP.

3. **One-Tailed Test**:
   A one-tailed t-test was used to assess whether the mean fluorescence of the fused genes was significantly higher than the unfused GFP. This approach focused on detecting whether the fused genes exhibited higher stability (i.e., greater fluorescence) than the unfused GFP.

4. **Calculation of P-Value**:
   The t-score was translated into a p-value using the standard t-distribution. The p-value was used to assess the statistical significance of the difference in stability between the fused genes and the unfused GFP. We calculated a p-value of 0.048. A p-value of less than 0.05 was considered statistically significant, indicating that the fused genes were significantly more stable than the unfused GFP.

5. **Interpretation of Results:**
   A significant p-value indicated that the fused genes had higher stability than the unfused GFP, supporting the hypothesis that gene coupling contributes to greater stability.

6. **Error Considerations**:
   Variability due to biological noise and experimental measurement errors was accounted for by incorporating the standard deviation of the fused genes and the standard error of the unfused GFP in the t-test calculation.

*Comparison between experiments*

We compared the stability of all experiments, to show that they do not have the same decay. The error analysis process was as follows:

1. **Data Collection**:
   For the 10 fused genes and unfused GFP, we collected all their fluorescence measurements along the experiment and normalized them by their fluorescence at time $t = 1$.
2. **Statistical test:**
   Under these assumptions, normality is no longer assumed. Rather, we performed a **Kruskal-Wallis H test**. This test does not assume normality, and has as a null hypothesis that all tests are equal. We derived a P-value of 0.035.
3. **Interpretation of Results:**
   A significant p-value indicated that the different fused genes and GFP come from different distributions, and do not have the same decay rate.
4. **Pairwise testing**
   Following this analysis, we conducted pairwise tests between the different fused genes and GFP, receiving significant results only for **SEC2**, with a p-value of 0.047. This indicates that top performing EG have a significant advantage over the baseline, while others may not – emphasizing the need for informed selection.

We further note that similar tests were conducted for other analyses:

- Conducting linear regression analysis on the log-values of the expression in the insulin experiment, we get that they are linear, indicating exponential decay ($P < 10^{-6}$). We also find that they have different decay rates ($ANOVA\ F - test, P < 10^{-19}$ ).
- By using the student's t-test for the leaky codon insulin experiment at time 40, we show that using a leaky stop codon provides slower decay than not using one ($P < 10^{-5}$) .
- By using the Kruskal-Wallis H test for the leaky codon insulin experiment, we show that each codon leads to a different expression profile ($P < 0.003$).

**Supplementary 2: Dataset Construction**

The dataset was taken from the SWAT library published in [1,2]. It contained fluorescence measurements of 6,685 endogenous genes in *S. cerevisiae*. They were engineered as fused genes, fused with either a NOP1 promoter and GFP or TEF with mCherry at the N' terminus of the gene. GFP and mCherry datasets were analyzed separately to account for any potential variation between labels. A correlation analysis revealed moderate alignment between GFP and RFP fluorescence.

All data were complete, and no imputation was required, as the features were derived directly from the genetic sequence. Fluorescence values were used as raw data without normalization to maintain their original scale.

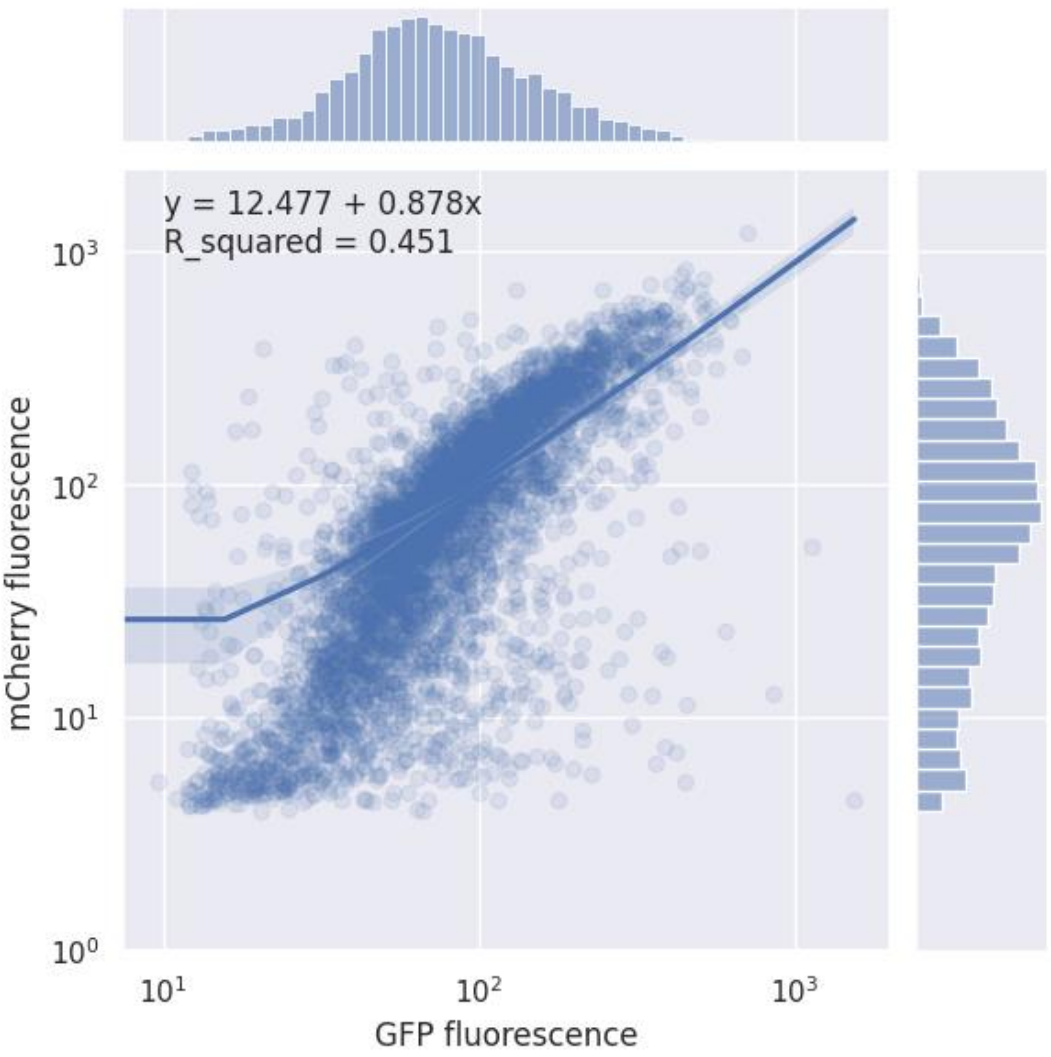


**Figure Sup. 1.1. Distribution and Relationship of Fluorescence Values for GFP and mCherry.** This graph illustrates the distribution of fluorescence values in the dataset for both univariate GFP and mCherry measurements, along with their relationship. Due to the wide range and skewed nature of the data, values are displayed on a logarithmic scale. A clear mutual trend is observed between GFP and mCherry fluorescence, as expected, indicating their correlated behavior in the dataset.

The data exhibited highly skewed labels, with correlations between the two target genes, GFP and RFP. To address this skewness, we employed a sub-sampling strategy that introduced variation across training sets, enabling the creation of an ensemble of models trained on different data subsets. This was repeated 20 times (see **Ensemble Approach**).

---

**Supplementary 3: Model Training and Evaluation**

**Ensemble Approach**

An ensemble approach was used, where 85% of the data was sampled for training in each iteration, and 15% held out for validation. This was repeated 20 times with different samples, to ensure robustness. Predictions were averaged among the ensemble models, for the final prediction.

**Models and Hyperparameters**

Four models were trained: XGBoost, KNN, SVR, and ElasticNet.

- **XGBoost**: Hyperparameters included learning rate (0.01–0.3), tree depth (3–10), subsample fraction (0.5–1), and regularization terms (lambda and alpha).[3]
- **KNN**: Number of neighbors (3–50) and distance metrics (Euclidean, Manhattan). [4]
- **SVR**: Kernel types (linear, RBF), penalty term (C = 0.1–10), and epsilon margin (0.001–0.1). [5,6]
- **ElasticNet**: Regularization strength (alpha = 0.01–1) and L1 ratio (0.1–1). [7]

Hyperparameters were tuned using Optuna, employing 5-fold cross-validation on training subsets.[8]

**Performance Metrics**

The **primary performance** metric was the median model result, expressed as quantile of the top fluorescence within test set in the top 3 candidates. Another performance metric measured was likelihood of failure, defined as probability of top fluorescence being below median performance within test set. These were also measured for performance for only top candidate, to check whether the model selected is consistent.

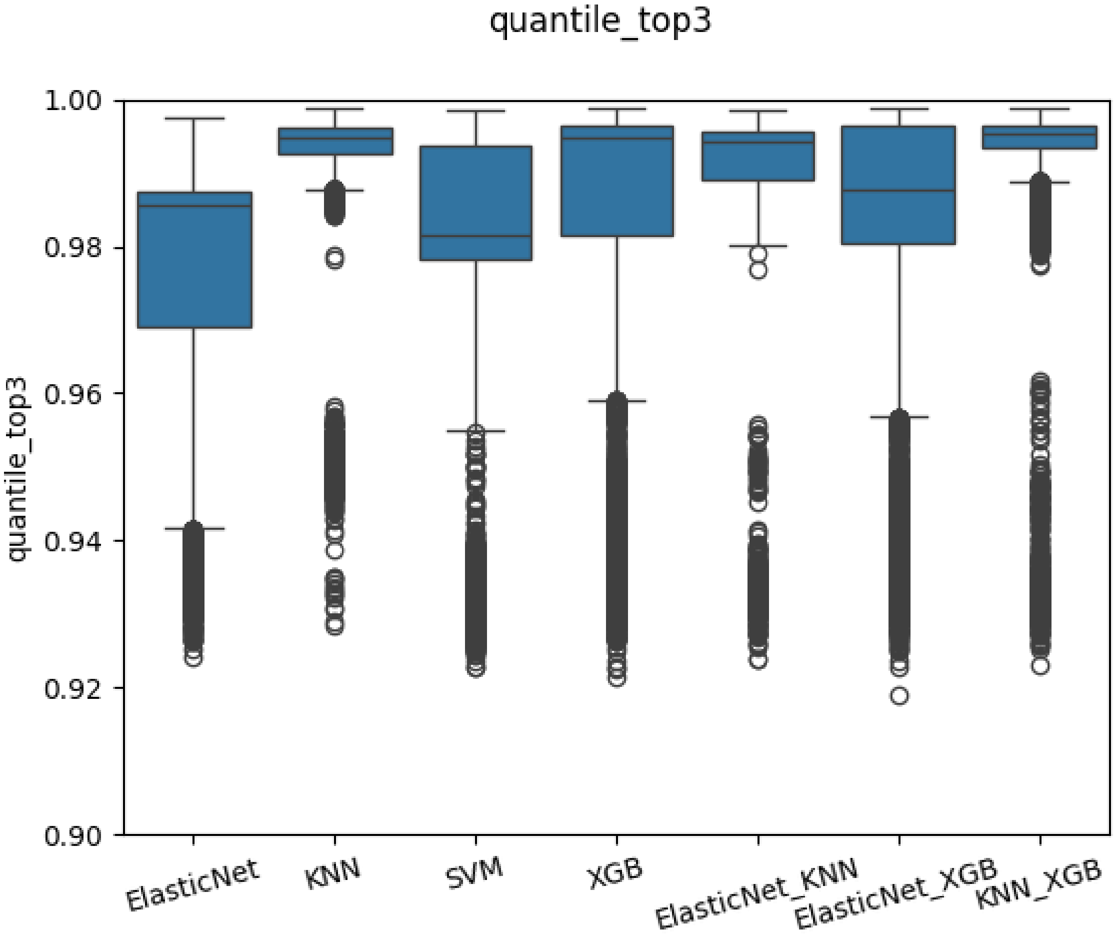


**Figure Sup. 3.1. Top Quantile of Actual Fluorescence Among Top 3 Candidates Across Model Architectures.** The graph shows the quantile of the top actual fluorescence (y-axis) among the top 3 predicted candidates for various model architectures. Each column on the x-axis represents results obtained by calculating the mean prediction for all models within a given architecture and applying the bootstrap method, where 6,685 genes were sampled with replacement (matching the original dataset size). The results demonstrate that model architectures incorporating K-Nearest Neighbors (KNN) exhibit a better capacity for catching highly-performing EG.

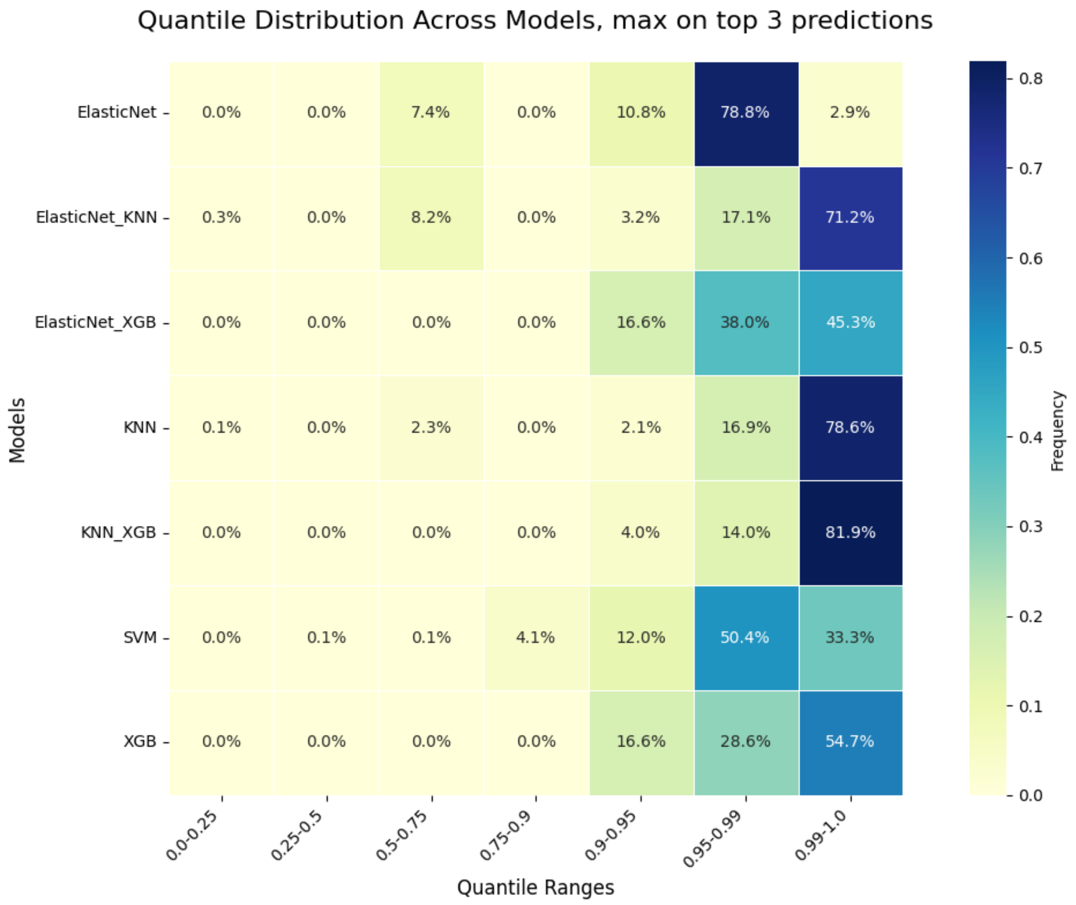


**Figure Sup. 3.2. Quantile Distribution of Actual Fluorescence Among Top 3 Candidates Across Model Architectures.** This heatmap displays the binned distribution of the quantile of top performing candidate among top 3 predicted. On one hand, ensembles involving KNN are more likely to find genes in top percentile. On the other hand, ensembles involving XGBoost are more robust, not getting quantiles under 0.9. The best results, both in likelihood to get top percentile and in robustness to failure, is an XGN and KNN ensemble.

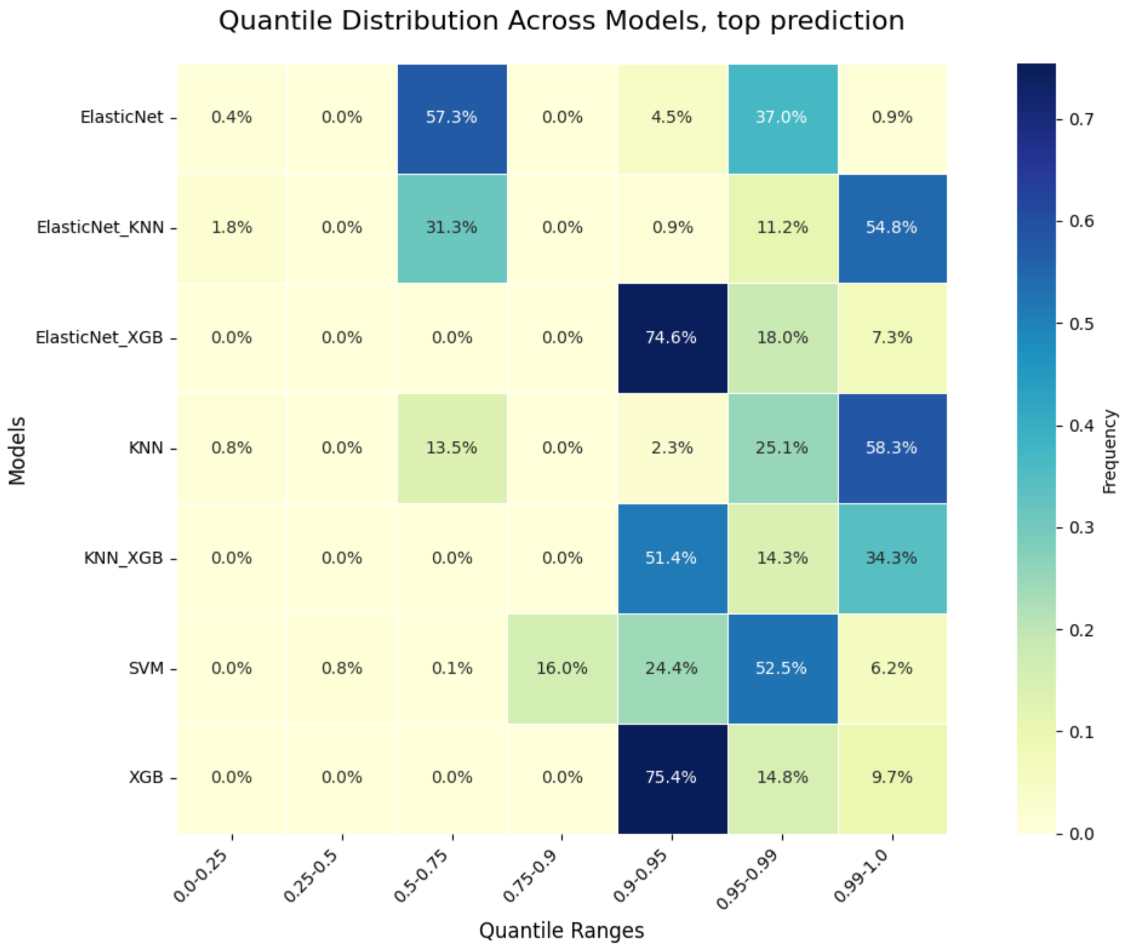


**Figure Sup. 3.3. Quantile Distribution of Actual Fluorescence of Top Candidate Across Model Architectures.** This heatmap displays the binned distribution of the quantile of predicted best candidate. The same pattern can be seen, where KNN ensembles are more likely to find top-performers, and XGBoost ensembles are robust to large errors. Thus, the XGBoost KNN ensemble displays the best performance, taking the best of both worlds.

Additionally, the models' performance was measured using the Spearman correlation [9], and AUC [10] for correctly classifying the top 5%. These metrics tested robustness (rather than performance on top candidates) and agree with the conclusion that ensembling KNN with XGBoost improved the model's performance.

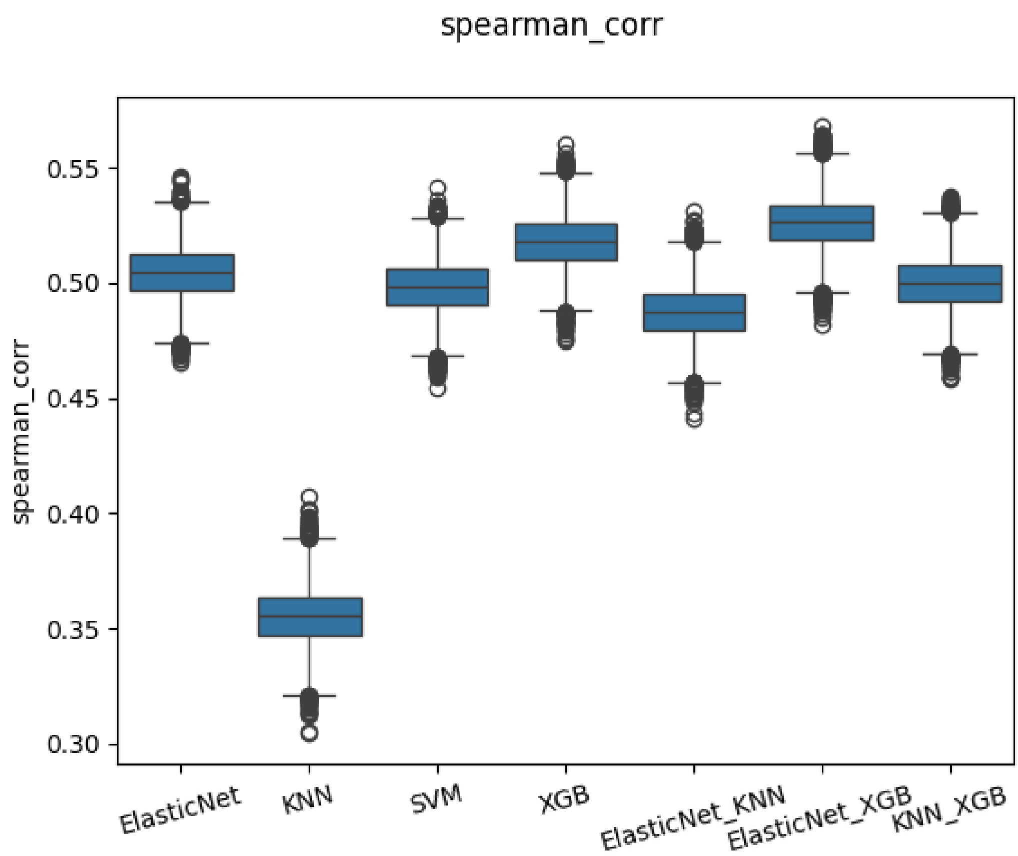


**Figure Sup. 3.4. Spearman Correlation between Prediction and Actual Expression Across Model Architectures.** This bar graph exhibits the distribution of Spearman correlation (y-axis)) between label and prediction for the different bootstrap samples. KNN exhibits lower robustness than the other architectures, further emphasizing the need for an ensemble model.

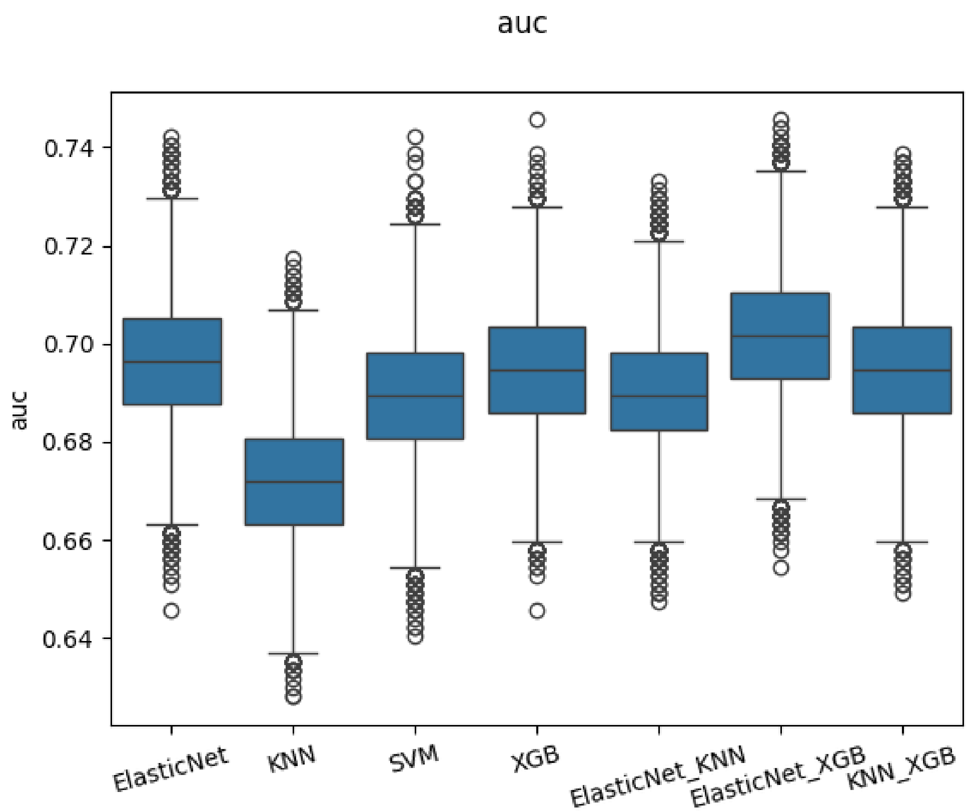


**Figure Sup. 3.5. ROC-AUC Performance for Top 5% Gene Classification Across Model Architectures.** The graph depicts the ROC-AUC (y-axis) for the classification task: "Is this gene in the top 5%?". Each column on the x-axis represents results obtained by calculating the mean prediction for all models within a given architecture and applying the bootstrap method, where 6,685 genes were sampled with replacement (matching the original dataset size). Among model architectures incorporating K-Nearest Neighbors (KNN), the combined KNN+XGB architecture demonstrates the most robust performance.

---

#### Feature Importance Analysis

Feature importance was derived for XGBoost using SHAP (SHapley Additive exPlanations), which ranks the contribution of each feature to model predictions. The top 20 features by SHAP values were used to assess the dominant predictors. Forward feature selection was applied to the KNN architecture, iteratively adding features based on their ability to maximize the AUC metric defined above[10].

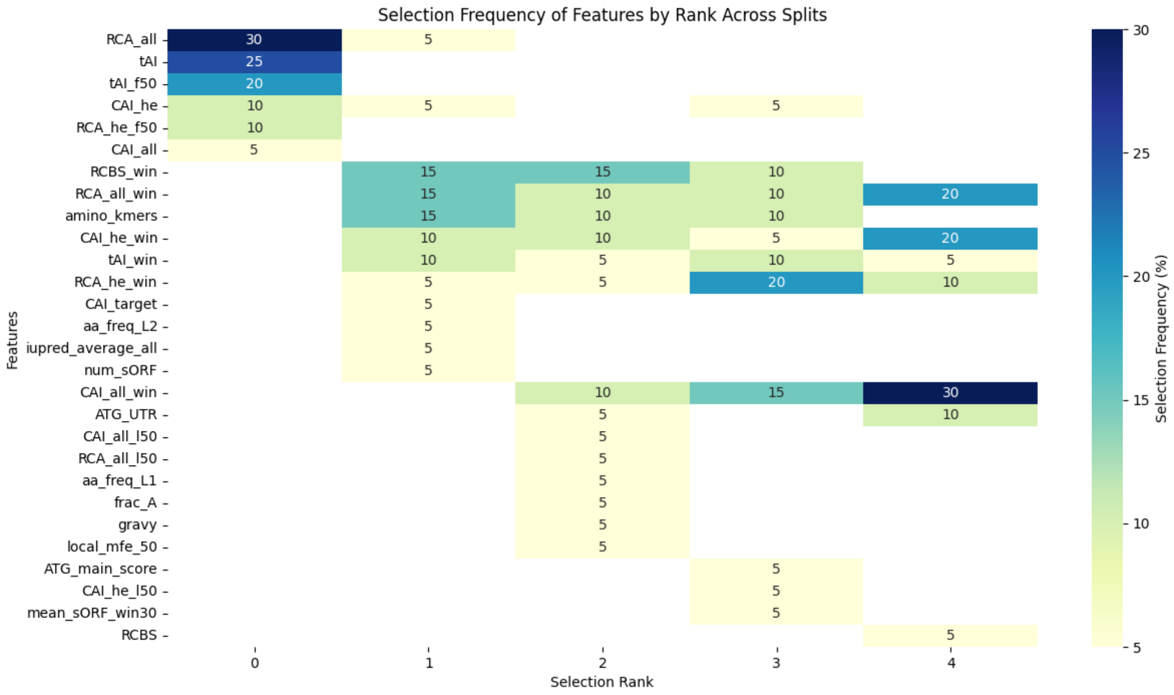


**Figure Sup. 3.6. Feature Selection Results from Greedy Forward Selection Across 20 KNN Models.** This graph presents the results of greedy forward selection for the top 5 features across 20 KNN models, where features were selected to maximize AUC, a robust performance metric. Each row represents a feature or family of features, and each column sums to 100% of selections for a specific selection rank.Features related to **Codon Usage Bias (CUB)** are the most frequently selected, with global CUB emerging as the most important, followed by CUB at initiation and downstream regions. Features related to **amino acid k-mer composition** are consistently selected in subsequent ranks. Other features contribute less significantly to the selection process.

### Supplementary 4: First Proinsulin Evolution Experiment, Linear Scale

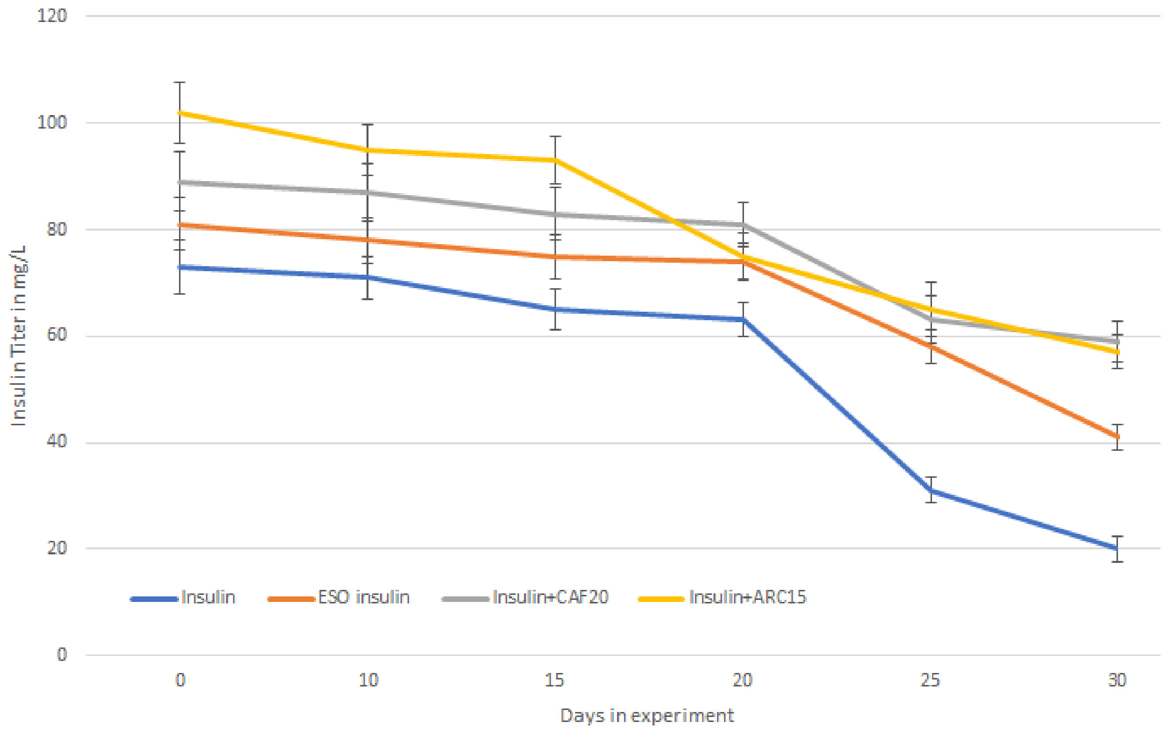


**Figure Sup. 4.1. Proinsulin production over time, Linear Scale.** This graph is a repeat of Fig. 3.C., presented in linear scale for further readability.
